# Response dynamics in macaque ventral stream recapitulate the visual hierarchy

**DOI:** 10.1101/2025.11.11.686115

**Authors:** Will Xiao, Kasper Vinken, Margaret Livingstone

## Abstract

Chronic recording in macaque inferotemporal cortex (IT) revealed a progression of feature selectivity over the duration of the response, with different images evoking strongest responses in different time bins. Neurons responded briskly and transiently to early best images and gave slower and more sustained responses to later best images. Correlations between IT selectivity and deep neural network (DNN) activations in early layers were faster and more transient than correlations with deeper DNN layers, suggesting a hierarchical representation unfolding. Analysis of best images in different time bins showed that early best images were simpler, or made up of fewer parts, than later best images. This simple-to-complex selectivity dynamic is consistent with, but more universal than, previously reported dynamics of global-to-local or category-to-exemplar selectivity. In parallel with the simple-to-complex selectivity progression, the image selectivity and the population activity became sparser during the response, consistent with the hypothesis of a canonical cortical microcircuit that pools inputs and incorporates neighboring-neuron inhibition. We propose that this temporal cascade in IT recapitulates the multiple antecedent stages in the visual hierarchy, which also show a simple-to-complex progression of feature selectivity.

## Introduction

Neuronal selectivity, especially in studies of the visual system, is often measured by summing spikes over a time window spanning tens to hundreds of milliseconds. This measure assumes homogeneous response selectivity over this summation window or, at least, that any changes in selectivity within this window are subordinate to the overall selectivity. The studies that have examined the dynamics of response selectivity invariably found that responses become more selective and sparser over the duration of the response, presumably because the effects of intracortical suppression evolve over this time. In macaque V1, about a third of the cells outside layer 4C become more sharply orientation selective over the course of the response. The earliest responses are broadly tuned to orientation, and become narrowed over time by broadly tuned intracortical inhibition (Ringach et al. 1997); see however (Gillespie et al. 2001). Most macaque V1 neurons and some IT cells show faster responses to low spatial frequency stimuli, and slower responses to higher spatial frequencies (Bredfeldt and Ringach 2002) (Toosti et al. 2024). Directional neurons in V1 and MT, when presented with a bar moving obliquely to its orientation, first respond to the motion perpendicular to the bar orientation, and only later represent the true motion direction, as intracortical suppression (end stopping) reduces the response to all but the ends of the bar (Pack and Born 2001; Pack et al. 2003). Posterior IT neurons respond faster to individual object parts than to multiple object parts (Brincat and Connor 2006). In central and anterior IT, the earliest response carries information about image category, with information about image identity arriving about 50ms later (Sugase et al. 1999; Tsao et al. 2006). In two face patches in anterior IT, response encoding of familiar and unfamiliar faces diverges only at long latencies (She et al. 2024). Lastly, anterior IT responses allow decoding of faces from non-faces earlier than decoding of individual faces from each other (Shi et al. 2023). Thus, in all these reports on the dynamics of visual responses, early spikes are less selective and less sparse than later ones. This convergent evidence across visual areas for sharpening selectivity over time suggests the possibility of a canonical mechanism underlying these diverse dynamics. Yet most previous studies have proposed task-specific, stimulus-specific, or area-specific explanations.

We hypothesize that a general principle of unfolding intracortical suppression governs the dynamics of visual selectivity. To test this principle across the visual hierarchy over a broad range of stimuli, we recorded chronically, up to weeks, from multi-site arrays in macaque IT and throughout the ventral visual pathway, while presenting large image sets up to 8.9k images. This allowed us to resolve unprecedentedly fine temporal dynamics and selectivity patterns. The results indicate a cascade over time of selectivity from simple to complex objects, mirroring an analogous progression along the visual hierarchy, and accompanied by increasing response sparsity across both images and neurons. This finding supports a universal mechanism of pooling followed by intracortical inhibition at each stage of the visual hierarchy, providing a parsimonious and general explanation for diverse previous reports of selectivity dynamics across visual cortex.

## Methods

### Animals

Seven male and two female adult macaques (3--14 kg) were used in this experiment; eight were rhesus macaques (Macaca mulatta) and one was a pigtailed macaque (Macaca nemestrina). They were implanted with custom-made titanium or plastic headposts and chronic microelectrode arrays in inferotemporal cortex, and, using long NHP Neuropixels probes, across occipital and temporal cortices (further details below). The monkeys were trained to perform a fixation task and were rewarded with juice for maintaining fixation on a spot in the middle of an LCD monitor 53 cm in front of the monkey. Gaze position was monitored using an ISCAN system (ISCAN, Woburn, MA). MonkeyLogic (https://monkeylogic.nimh.nih.gov/) was used as the experimental control software. All procedures were approved by the Harvard Medical School Institutional Animal Care and Use Committee (protocol #ISO00001049) and conformed to NIH guidelines provided in the Guide for the Care and Use of Laboratory Animals.

### Visual Stimuli

Images included objects on a white background from either (Konkle and Oliva 2012) or (Brady et al. 2008), comprising several hundred categories of objects with 1–17 exemplars per category, or photographs taken by M. Livingstone of monkeys, humans, and objects in our lab, familiar to the monkeys (up to 180 images per category). In total, our image sets contained 177 to 436 categories, with an average of 4 to 28 images per category (for example, cookies, knives, cooking pots, vases, big animals, small animals), depending on the overall image-set size. We used increasingly large image sets over the duration of the study. For monkey R, the data in Figures 1&3 were collected using a 776 image set, whereas monkey R data in subsequent figures were collected using a larger image set. While the monkey maintained fixation, images were presented at a size of 4–6 visual degrees at the center of the array’s mapped receptive field. We used different image presentation times for different monkeys depending on the response time-course for each array. Among the data sets in this study, each image was presented for 9 to 59 repetitions (Supplementary Table 1).

**Figure 1.**
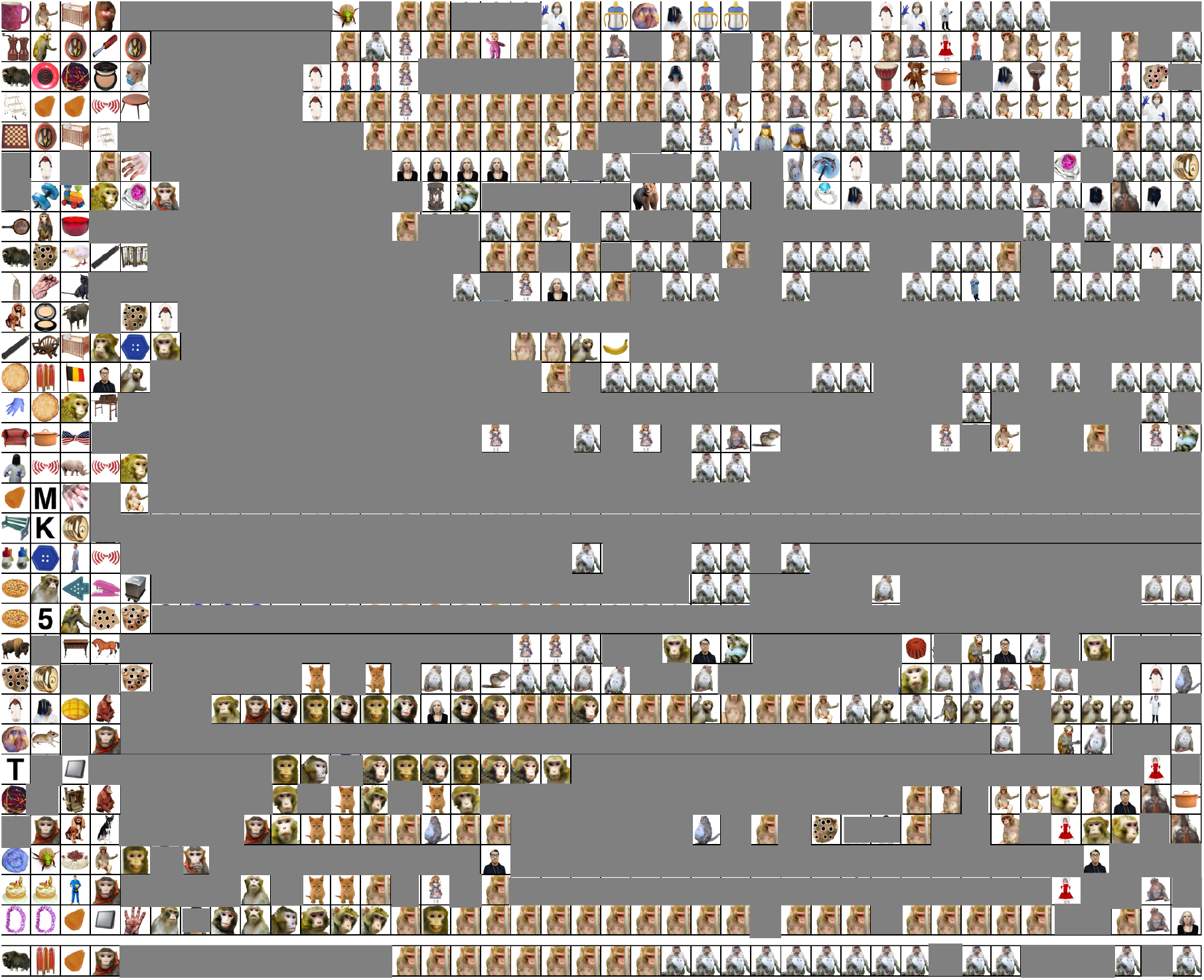
Response dynamics from a NiCr microwire array in AIT of monkey R showing shifting image selectivity. The figure shows the top image for each of 31 recording sites (rows) for each 10ms time bin from 60 to 450 ms after stimulus onset; all bin responses were at least 2 standard deviations above baseline. The last row corresponds to responses averaged over the array. The most common images in the image set (67%) were inanimate objects, of which reddish rounded objects account for only 15% of the full image set. All images of humans have been grayed out, except for photographs of authors; the remaining human-like figures are inanimate dolls.

**Figure 2.**
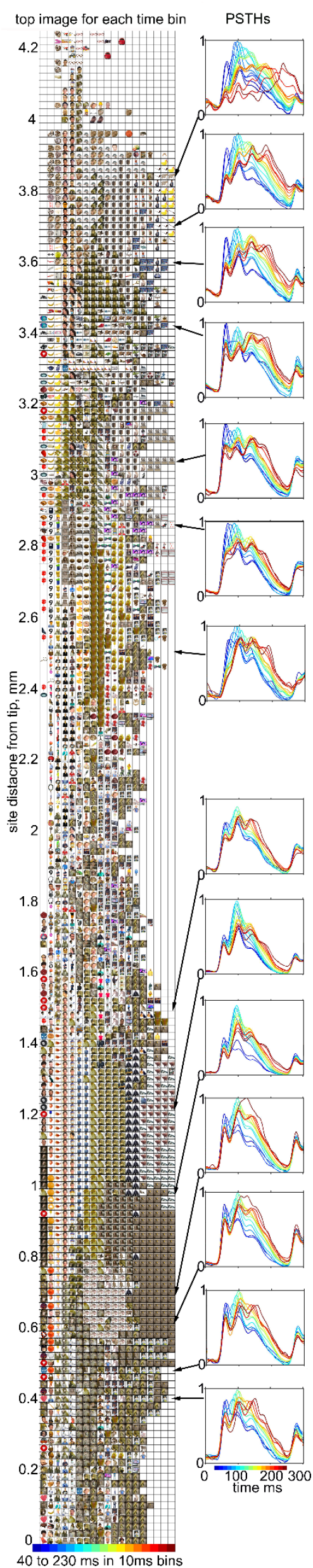
Response selectivity dynamics from a Neuropixels recording in the middle face patches of monkey B5. (left) Top images for each 10ms time bin (columns) from 40 to 230 ms after stimulus onset for 212 recording sites (rows) from a chronically implanted 10mm Neuropixels probe in the CIT face areas of monkey B5. Images are plotted only for responses at least 2 standard deviations above baseline. (right) PSTHs (normalized to the maximum response across time bins) in response to the top 100 images from each 10ms time-bin (color bar on the time axis). Magnified images for example sites are shown in Supplementary Figure 5.

**Figure 3.**
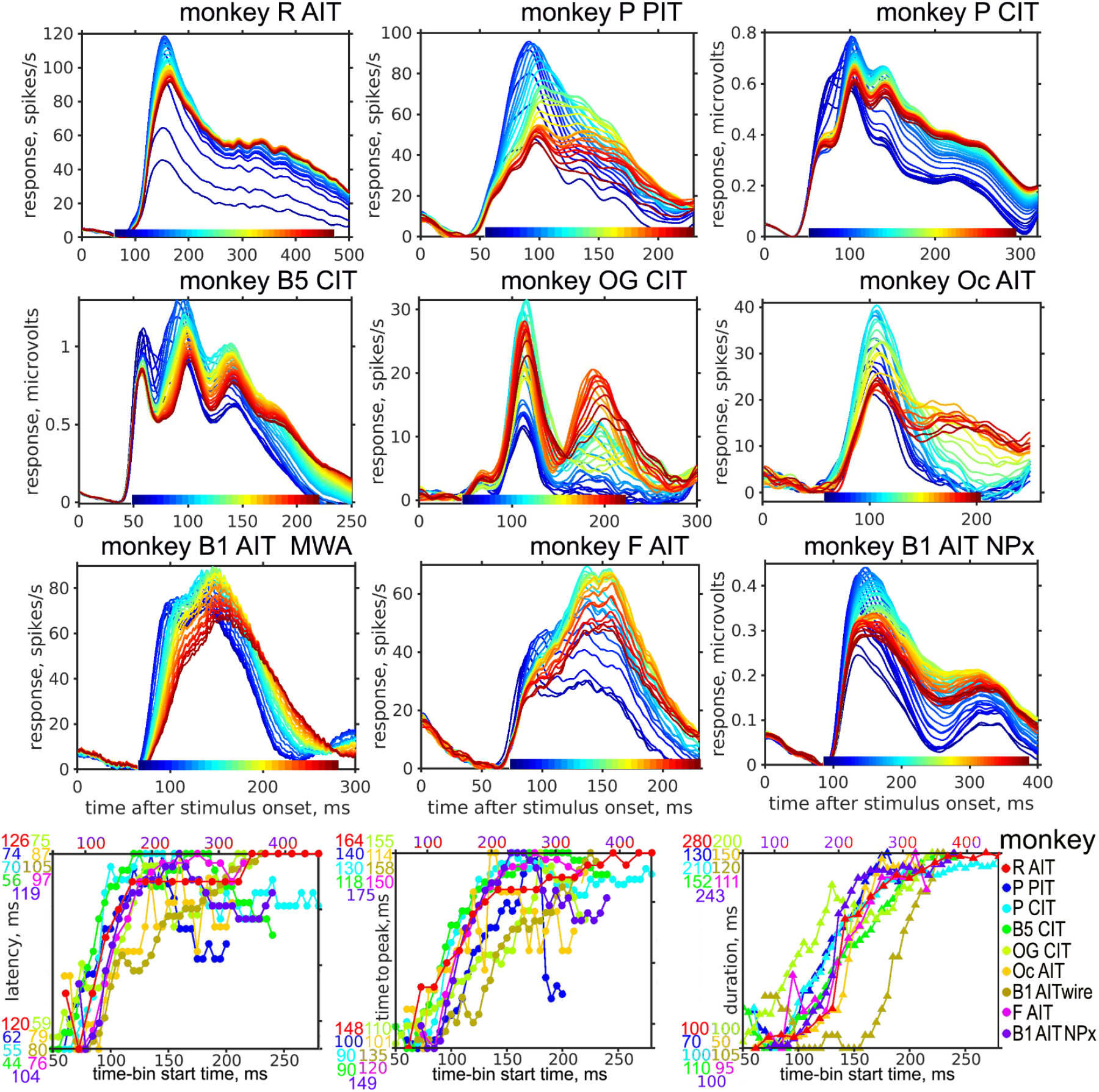
(top 3 rows) PSTHs for early top images (blues) are faster and more transient than PSTHs in response to later best images (reds). The upper 9 graphs plot the array-average PSTHs in response to the top 100 images from each 5ms time-bin (10ms for monkey R) spanning the response duration. For each time-bin indicated by color along the time axis, responses to the top 100 images are plotted in the corresponding color. Early best-image responses are faster and more transient than responses to later time-bin best images. (bottom row) Summary response dynamics for 9 arrays in 7 monkeys. The array-average PSTH latency, peak time, and duration for each array are plotted with the array indicated by color. The y-axis ranges are normalized per array, with the maximum and minimum values per array indicated by colored numbers so that the actual values can be ascertained. The upper x-axis range corresponds to datasets from monkeys R and B1-AIT-NPx, which had slower responses and longer stimulus presentation times. The lower x-axis range corresponds to the other datasets.

### Physiological recording

Animals were chronically implanted with a custom floating microelectrode array (FMA, 32 channels, MicroProbes, Gaithersburg, MD), a microwire array (MWA, 64 channels; MicroProbes), or a short (10mm) or long (45mm) Neuropixels array (IMEC, Leuven, Belgium). Each animal received 1–3 arrays throughout data collection spanning three years. The arrays were sealed to the skull with surgical silicone and dental cement. We recorded from 13 chronically implanted arrays in 9 monkeys. One array was an FMA Floating Microelectrode Array (Microprobes for Life Sciences) implanted in the posterior face patch in IT; individual recording sites were separated by 400 microns, and the maximum distance between sites was 3.5mm. Five arrays were MWAs, which are bundles of fine (12μm diameter) wires that splay out and float in the cortex. The wire tips remained within 1–2mm from each other as assessed by high resolution cone-beam CT (i-CAT FLX-V, Dexis, Quakertown, PA); Supplementary Figure 1; see also (McMahon et al. 2014). Two of the arrays were 10mm long Neuropixels probes and five were 45mm long Neuropixels probes (IMEC, Leuven, Belgium). The data for monkey B1 AIT NPx was collected from the tipmost 7.7 mm of a 45mm probe, in order to have a homogeneous IT population. Of the MWAs and Neuropixels arrays, some were targeted to face patches using fMRI and anatomical markers to identify face selective regions (Arcaro and Livingstone 2021), whereas other arrays were outside face patches and not face-selective (Supplementary Figure 3).

Neural signals from FMAs and MWAs were amplified and sampled at 40 kHz using OmniPlex data acquisition systems (Plexon, Dallas, TX). Multi-unit spiking activity was detected online using a threshold-crossing criterion at 3.5 standard deviations. Channels containing separable waveforms were sorted online using a template-matching algorithm. Some sites had a few well-isolated single units that were stable over the duration of recording, while most of the data were multi units. There was no qualitative difference in the dynamics of the single units from the multi-units. Neuropixels data were acquired at 30 kHz (action-potential band) using SpikeGLX software (Janelia Research) and preprocessed using CatGT, which includes global-average referencing. We quantified activity by the multi-unit activity envelope (MUAe). The MUAe was calculated with custom code that, following previously described methods (Super and Roelfsema 2005; Chen et al. 2020), performed bandpass filtering (forward-backward Butterworth filter, effectively 6^th^ order, 300–3000 Hz), rectification (taking the absolute value), lowpass filtering (at 200 Hz), and finally down-sampling to 1 kHz. The resulting MUAe provides an instantaneous measure of aggregate spiking activity near the electrode (Legatt et al. 1980; Brosch et al. 1995) and does not depend on an arbitrary spike-detection threshold. For all arrays, neural signals were synchronized by TTL events to task data. Synchronized image-onset event times were further refined using photodiode signals.

### Stable Chronic Recording

The chronically implanted arrays provided stable recording for weeks, even months, allowing us to present hundreds to thousands of images with 9–59 repetitions over recording sessions (Supplementary Table 1). We assessed recording stability for the datasets used in this study by the day-to-day consistency of image selectivity at the categorical and individual-image levels (Supplementary Figure 2).

### FMA and MWA data registration across sessions

We identified multi-units recorded on each electrode with activity on the same electrode over days. In cases there were sorted units, they were always stable, which we confirmed daily during data acquisition and online sorting using waveform templates saved from the previous day.

### Neuropixels data registration across sessions

Chronic Neuropixels data were registered using rigid 1D shifts based on the trial-averaged evoked MUAe response (analogous to PSTHs), a matrix of shape time x neurons. Thus, the registration used the spatiotemporal profile of responses along the probe but not any image selectivity. We arbitrarily selected a session as the anchor session and then registered all other sessions to this one. Each moving session was aligned independently to the anchor session. We searched a grid of shift values (−500 µm to 500 µm in 1 µm steps, finer than the spacing between electrodes because we used interpolation to optimize a single shift parameter, pooling information across electrodes). With each shift value, we shifted the nominal electrode positions of the moving session, 1D-interpolated its evoked response matrix to the electrode positions of the anchor session, calculated the correlation between response matrices, then selected the shift value that maximized this correlation. This procedure resulted in a typical shift value of tens of microns. To downweigh nonresponsive and noisy sites, we used temporal smoothness as a prior, scaling the evoked response of each site proportional to how similar (R^2^ score) the evoked response was to its smoothed version (25 ms moving window averaging); the R^2^ score was normalized to a range of 0–1 based on the 2.5– 97.5^th^ percentile across electrodes, then multiplied onto the responses per electrode, thereby weighing how much each electrode contributes to the optimization of the between-session alignment. In one case (monkey P’s long NPx probe; Figure 7) where we configured to record from disjoint sections of the probe, we separately aligned each contiguous section of electrodes. Although a more complex registration algorithm could be used (e.g., as in Kilosort 2.5 (Steinmetz et al. 2021), fitting a spline interpolation with multiple probe sections, allowing the registration to vary within a session, and using, instead of a fixed anchor session, a template iteratively optimized with expectation-maximization), we elected to use a simple algorithm, thus providing a conservative estimate of the chronic stability of recordings.

After all sessions were registered, data were concatenated by applying the optimized shift to every moving session and then interpolating the responses onto the site locations of the anchor session. Because the MUAe magnitude is sensitive to day-to-day fluctuations in the raw signal magnitude, we scaled the responses of every moving session to match the overall background activity and peak response magnitude of the anchor session, using two parameters (a slope and an intercept) per session per electrode. To calculate these parameters, we first smoothed the evoked responses with a 50-ms moving average window and then downsampled the responses to 100 Hz. The background activity was calculated as the median activity (across presentations and time bins) between −100 and 0 ms relative to image onset. The peak response magnitude was calculated as max (across time bins) median absolute deviation (across presentations) of the response between 50 and 500 ms post image onset. The slope and intercept were chosen to match the background and peak activity from the moving session to the anchor session.

### Selectivity stability (Supplementary Figure 2)

We calculated selectivity correlation between days at the category and individual-image levels. To calculate category selectivity correlation between days, we randomly split (13 times) each day’s presentations into halves, stratifying by image category, and then correlated each half of one day with each half of the other day, resulting in 13 × 2 × 2 = 52 correlation values per day pair. The 52 correlation values were averaged. To calculate category selectivity correlation within a day, i.e., self-consistency at the category level, we randomly split (100 times) the presentations within a day into halves, again stratifying by image category, and correlated the two halves. To ensure the quantification included only visually selective neurons, we kept only neurons with within-day self-consistency ≥ 0.4, using the average of the first 50 of 100 within-day splits to select neurons and reporting the average of the second 50 splits. Analogously, to select visually selective neurons for between-day quantifications, we use the within-day self-consistency of a random day from the pair of days to select the neurons. Therefore, the between-day selectivity stability is a more conservative estimate than the within-day self-consistency. The same quantification was repeated at the individual-image level (bottom plots).

### Quantification and Statistical Analyses

We included only visually responsive sites, defined by having evoked responses at least 2 standard deviations above baseline. All PSTHs were cross validated by taking the top images for each time-bin from the first half of recordings, and averaging responses to those same images from the second half of recordings, and vice versa, then averaged. Virtually identical results were obtained by splitting the responses into odd and even trials.

### Representational similarity analysis (Figures 5 & 8)

The neuronal population representation of an image was defined as the vector of responses across neurons, with responses normalized per neuron across images. An image-image pairwise distance matrix was calculated using cosine distance (one minus the normalized dot product between image representations). An analogous pairwise distance matrix was calculated for features in deep neural networks (DNNs).

Representational similarity was calculated as the correlation between the two distance matrices, whereof we used only the upper-triangular part of each distance matrix because the matrix is symmetrical with constant zeros in the diagonal. Different response time-bins were analyzed independently, as were different model layers.

For the DNN, we analyzed pre-trained AlexNet (Krizhevsky et al. 2012), ResNet-50 (He et al. 2016), and ViT-L/16 384 × 384 (Dosovitskiy and Brox 2016). We selected layers roughly evenly spaced in depth, except for AlexNet where we analyzed all eight layers (ResNet-50: 16 selected layers; ViT: nine selected layers). Each model feature (hereafter, ‘unit’) was averaged over the width and height dimensions (convolutional layers in AlexNet and ResNet-50) or the sequence dimension (ViT). AlexNet layers had units ranging from 64 D to 4096 D, the selected ResNet-50 layers had units ranging from 256 D to 2048 D, and all selected ViT layers had units of 1024 D.

For spatial resolution along a Neuropixels probe (Figure 7), we grouped the sites into groups of 10 adjacent sites (advancing by 5 sites) and then analyzed each group as a population. To visualize the vector of RSA fit values across model layers as one cell in a heatmap, we determined the weighted best-fit layer (relative depth) as follows. The RSA fit was normalized between 0 (min across layers) and 1 (max) and then softmax-transformed into a weight vector (i.e., sums to 1), with the temperature hyperparameter set to 0.2 (i.e., an increase of 0.2 in normalized RSA fit increases relative weight by about 2.7-fold; as temperature approaches 0, the softmax function reduces to the max function). This weight vector was used to calculate the weighted-average relative depth across layers:

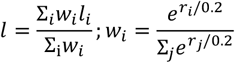

Where *i* indexes the layers and *l*_*i*_ = ∈ (0, 1] is the relative depth of each layer. Using the simple max to visualize the best-fit layer depth leads to the same conceptual conclusions, while the weighted depth shows more value gradations.

### Image complexity

Image complexity was quantified by how many parts each image consisted of by first clustering pixels by color using k-means clustering (k=4) (Späth 1985) and then counting the number of unconnected components in each color cluster.

### Image spatial frequency

The average spatial frequency of each image was estimated as follows. We first converted color images to luminance (gray scale), applied a two-dimensional fast Fourier transform (fft2), and centered the zero-frequency component with fftshift. Then the power spectral density (PSD) was obtained as the squared magnitude of the Fourier coefficients, *P*(*u, v*) = |*F*(*u, v*)|^2^, where u and v index the x and y spatial frequency components. The (orientation-agnostic) spatial frequency was computed as 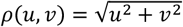 (cycles per image). The image’s average spatial frequency was defined as the power-weighted mean spatial frequency:

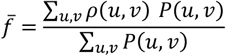

Because *P*(*u, v*) integrates to the total signal power, this measure is invariant to absolute image contrast.

### Image sparsity (sometimes referred to as ‘lifetime sparsity’)

For each time-bin, for each visually responsive site, the top 5% of the images were found using cross-validated responses. To do this, the image rank order based on image responses averaged over even stimulus presentations was used to sort image responses averaged over odd stimulus presentations, and vice versa. These image-responses were then robustly normalized per site per time-bin between the 2.5^th^ and 97.5^th^ percentiles. The normalized responses for each image were used to calculate image sparsity using the following formula (Treves and Rolls 1991):

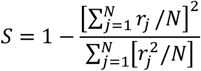

where r_j_ is the response to the j^th^ image. We report S averaged over all visually responsive sites for each array.

### Neuronal population sparsity

For each time-bin, for each visually responsive site, the top 5% of the images were found as above. Then, for each image, the normalized responses (normalized per site across images as above) were used to calculate sparsity with the same formula above, except r_j_ is the response of the j^th^ site out of N visually responsive sites. We report S averaged over the top images.

### Interpolation to a normalized time axis (Figure 6)

To plot the varied time-courses for all 9 focal arrays on the same time axis (Figure 6), we interpolated each array time-course between the two troughs in the PSTH, as indicated in Figure 4.

## Results

We recorded from 9 focal chronic arrays in 7 monkeys. Some of the arrays were in face patches and were face selective, while others were not (Supplementary Figure 3. The recording sites in each array tended to have similar face selectivity, presumably due to the small spatial spread of electrodes in each array.

### Dynamic shifts in image selectivity in IT

We initially noticed dynamic shifts in image selectivity when we plotted the images giving the strongest responses as a function of time after stimulus onset. To illustrate this, Figure 1 shows the time-evolving top image (the image with the highest average response) for each of 31 electrodes, as well as the array average, of a microwire array chronically implanted in a face patch in anterior IT of monkey R (same array as in Supplementary Figure 1 left; red dots in Supplementary Figure 3). Each column corresponds to a 10ms time-bin from 60 to 450 ms after stimulus onset. The typical face selectivity index of sites in this array was 0.87 ± 0.2 (mean ± s.d.), and all the individual recording sites in this array had a face-selectivity-index above 0.3. Over 13 days of recording, 776 color images on a white background were presented for 200ms ON/200ms OFF at 5 degrees across, centered on the population receptive field of the array. By inspection, most top images depict people wearing PPE (despite only 7.7% of the image set being people wearing PPE) or monkeys (8.4% of the image set). Most neurons showed face preference only around 100 ms after stimulus onset. The earliest best images were reddish rounded objects (15% of the image set), not faces. From around 150 ms, the best images tended to be torsos, and later whole individuals.

There is a shift over time in the category of the top image for this example array (Supplementary Figure 4a). The most common category among the top images gradually shifted from reddish rounded objects to faces, torsos, and finally to whole individuals. This shift was evident despite the stable ranking of category-average responses, which showed that faces on average were preferred throughout the response duration (Supplementary Figure 4b&c). The shift in the category of the top image is also reflected in different PSTH time-courses of the categories. Reddish rounded objects evoked the fastest, most transient responses, followed by other objects, then faces, and finally whole humans or whole monkeys, which evoked the slowest, most sustained responses (Supplementary Figure 4 d&e). The response to reddish objects is larger than the response to all other inanimate objects only very early in the response, when the firing rate above baseline was low. Nonetheless, the difference is significant: the response across recording sites between 75 to 100 ms after stimulus onset to reddish objects (1.1 spikes/s above baseline) is significantly larger (2-tailed ttest, p=5.8 *10^-4^) than the response to all other inanimate objects (mean across sites = −0.2 spikes/s).

It is difficult to interpret this shifting selectivity over time as an evolution towards the optimum stimulus. The top image changed continually, and faces were the top image only transitorily, yielding to torsos and whole individuals later in the response. The distinct preferred images at different phases of the dynamic response exhibited different (transient vs sustained) response time-courses.

Selectivity changed over the response duration in IT across arrays and monkeys. For another example, Figure 2 shows the top images per 10 ms time bin for 212 electrodes on a short (10mm) Neuropixels probe chronically implanted in CIT of monkey B5. This array intersected the lateral middle face patch (depths 4.2 to 2.2 mm) and the fundal middle face patch (depth 1.6 mm to tip). The image set consisted of 3212 images on a white background, half of which were inanimate objects. Images were presented 100ms ON/ 100ms OFF. Without discerning each image, the changing selectivity over time is nonetheless apparent; larger images for eight example sites are shown in Supplementary Figure 5. The dynamics were similar to those in the first example array (Figure 1): Most of the earliest top images are round warm-colored objects, followed often by faces, then by torsos or whole humans, monkeys, or other whole mammals.

We hypothesize that these response dynamics can be explained by canonical principles, namely hierarchical pooling and intracortical feedback, that are general across visual cortex. To avoid relying on categories to explore principles underlying response dynamics, we instead grouped the top images in each time-bin and compared the average PSTHs in response to these different groups of images. For brevity, we refer to top images in early (late) time bins as early (late) best images. By inspection, as with monkey R, all the example sites in the B5 array showed fast transient responses to the early best images (blues) and slower more sustained responses to the later best images (reds) (Figure 2, right column). Early best images, which produce strong transient responses, are no longer the best images later in the response. While early best images are expected to evoke faster responses, it is not foregone that these responses should be more transient than responses to late best images. Nor is it expected that some images would produce fast, transient responses and others slower sustained responses. A shift of preferred images from those that elicited fast, transient responses to images that gave slower sustained responses was observed in all 9 focal arrays, including arrays that were not face selective (Figure 3). This shift was observed for both individual recording sites and array-average responses.

We quantified the response dynamics of top images from different time-bins by finding the time when the response reached half the peak magnitude (the latency), the time of the peak response, and the time between the latency and the first time point when the response returned to half the peak (the duration). Figure 3 (bottom) summarizes the latency, peak-time, and duration of responses to the top-100 (top 50 for monkey R) images per time-bin for 9 arrays in 7 monkeys. Responses tend to have a shorter latency, faster time to peak, and shorter duration for early best images and slower and more sustained responses for late best images.

### DNN representational similarity shifts from shallow to deep layers over the response duration

Given the various previously reported observations about response dynamics in IT: local→global, category→identity, few parts→many, low spatial frequency→high, average face→unique faces, coarse→fine, and unfamiliar→familiar, we asked whether some common principle might be revealed by comparing neuronal selectivity dynamics to the activations in different DNN layers. It has been shown that later layers of DNNs predict macaque IT responses, whereas V4 responses are best predicted by middle layers (Yamins et al. 2014). We calculated the image-image pairwise distance matrix for each neuronal population, and a second pairwise distance matrix for DNN activations. We tested three DNNs, AlexNet, ResNet-50, and ViT-L/16 384 × 384 (Krizhevsky et al. 2012; He et al. 2016; Dosovitskiy et al. 2021). The representational similarity between the neuronal and DNN distance matrices was calculated as a function of time (in 5ms time-bins) for each selected DNN layer. Figure 4 shows the dynamic RSAs with ViT-L for each array population. For both ViT-L and AlexNet, although not ResNet-50, the RSAs for shallow layers (blues) tended to rise earlier than for deeper layers (reds) for most of the arrays (Figure 4 top). To quantify this we plotted the latency (time to half peak) of the RSAs with different DNN layers (Figure 4 bottom row). For all 3 networks representational similarity with shallower DNN layers generally had shorter latencies than representational similarity with deeper layers. The 3 shallowest layers in each DNN had shorter latencies than the deepest 3 layers across arrays (2-tailed t-test, Alexnet p = 2.7 x 10-5; ViT-L p = 4.1 x 10-6; ResNet p = 0.48). The poorer latency correlation with ResNet-50 layers may reflect the less distinct representations among layers in deep ResNets (Kornblith et al. 2019).

**Figure 4.**
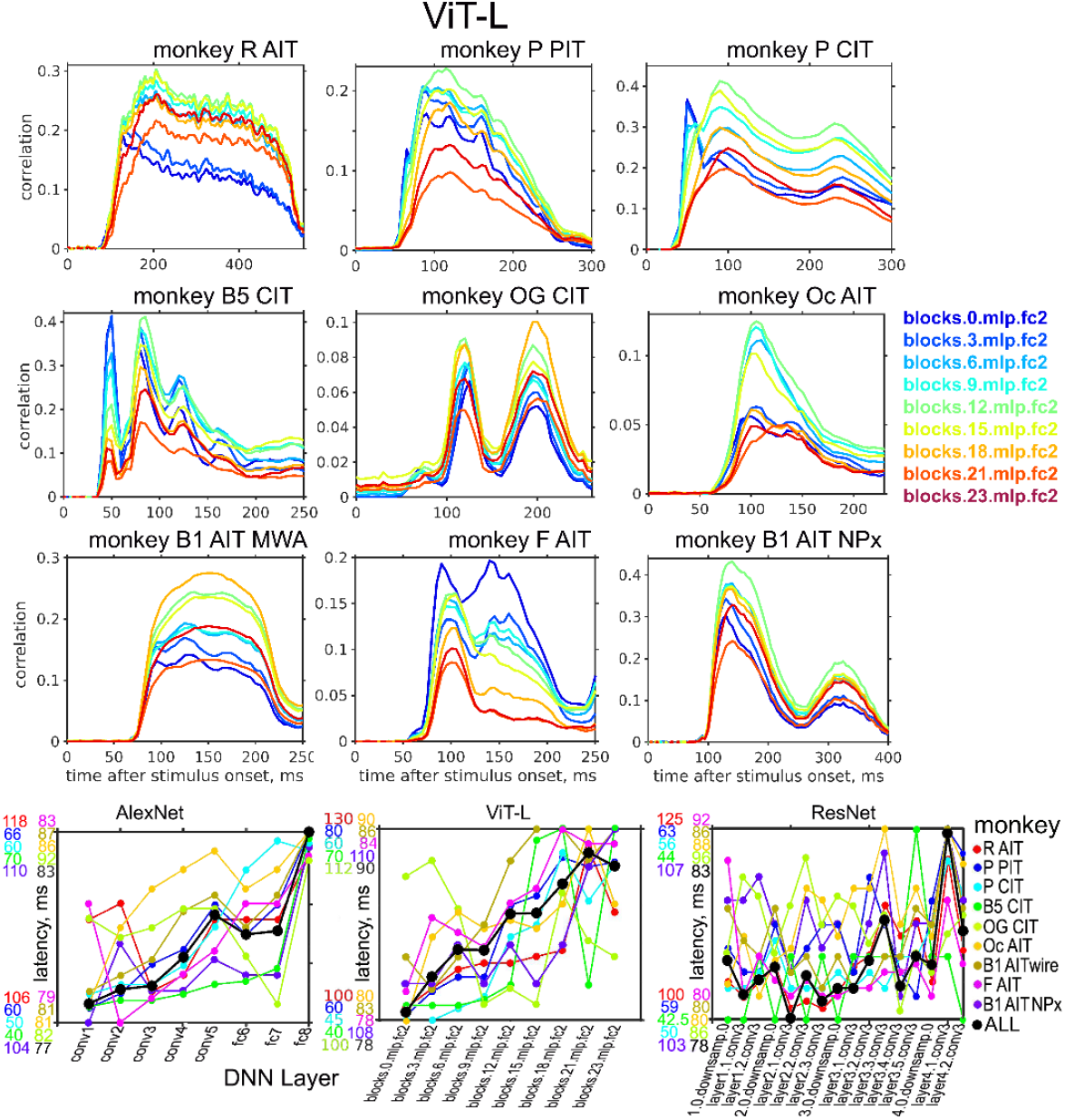
Representational Similarity Analysis (RSA) (Nili et al. 2014) between neuronal populations and different layers in the deep neural network ViT-L. The upper 9 graphs plot, for each array, the time-varying RSA (Pearson’s correlation) between the neuronal population responses in each time-bin and the activations in the indicated layers of ViT-L. For most of the arrays RSAs peaked earlier for shallower layers (blues) and later for deeper layers (reds). The bottom graphs plot the latency of the RSA time-course of each array (indicated by color; black indicates average over all arrays) with three different DNNs. The y-axis ranges for each monkey are indicated by colored numbers.

Feature visualizations of DNN units show that those in shallow layers encode ‘simple’ features, like contours or Gabors, while those in deep layers encode more ‘complex’ feature conjunctions that map well onto faces or entire objects (Olah et al. 2017). Deep neural networks approximate compositional functions better than shallow networks (Poggio et al. 2017), and deeper layers show increasingly better classification accuracy thanks to increasingly separable object representations (Cohen et al. 2020). The output layer of Therefore, the finding that neuronal correlations with DNN layers are faster for shallower than deeper layers is consistent with the idea that early responses in IT are more selective for simple images (represented by shallower layers), while later parts of the response are more selective for complex images (deeper layers).

### Increasing preferred-image complexity over the response duration

Instead of DNN representations, a simpler and more direct measure of object complexity is its number of parts (most of our images were of single objects on a white background). The number of object parts is also germane to the hypothesis that pooling operations at each stage combine features from prior stages (Riesenhuber and Poggio 2000). We therefore quantified how many parts each image consisted of by simply counting the number of non-contiguous color clusters. Because the image segmentation algorithm counts contours as separate clusters, it overestimates what we would intuitively label as image parts, yet it consistently quantifies more complex images as having more parts. Supplementary Figure 6 shows the segmentation of a few example images into parts.

The parts-based complexity measure confirms that the earliest IT responses are selective for images with fewer parts, while later responses prefer images with more parts (Figure 5). For all the arrays except monkey R’s, before the neuron starts responding to an image presentation, the average preferred-image complexity is similar to the average complexity over the entire image set (red dashed line), as expected, since the top images before response onset should be a random subset of the images. The above-average complexity early in monkey R’s plot probably reflects carry-over from the preceding stimulus. As the cells start responding, the mean complexity of the top 100 images per time-bin plummets significantly below the image-set average, then rises gradually over the duration of the response to significantly above the image-set average.

**Figure 5.**
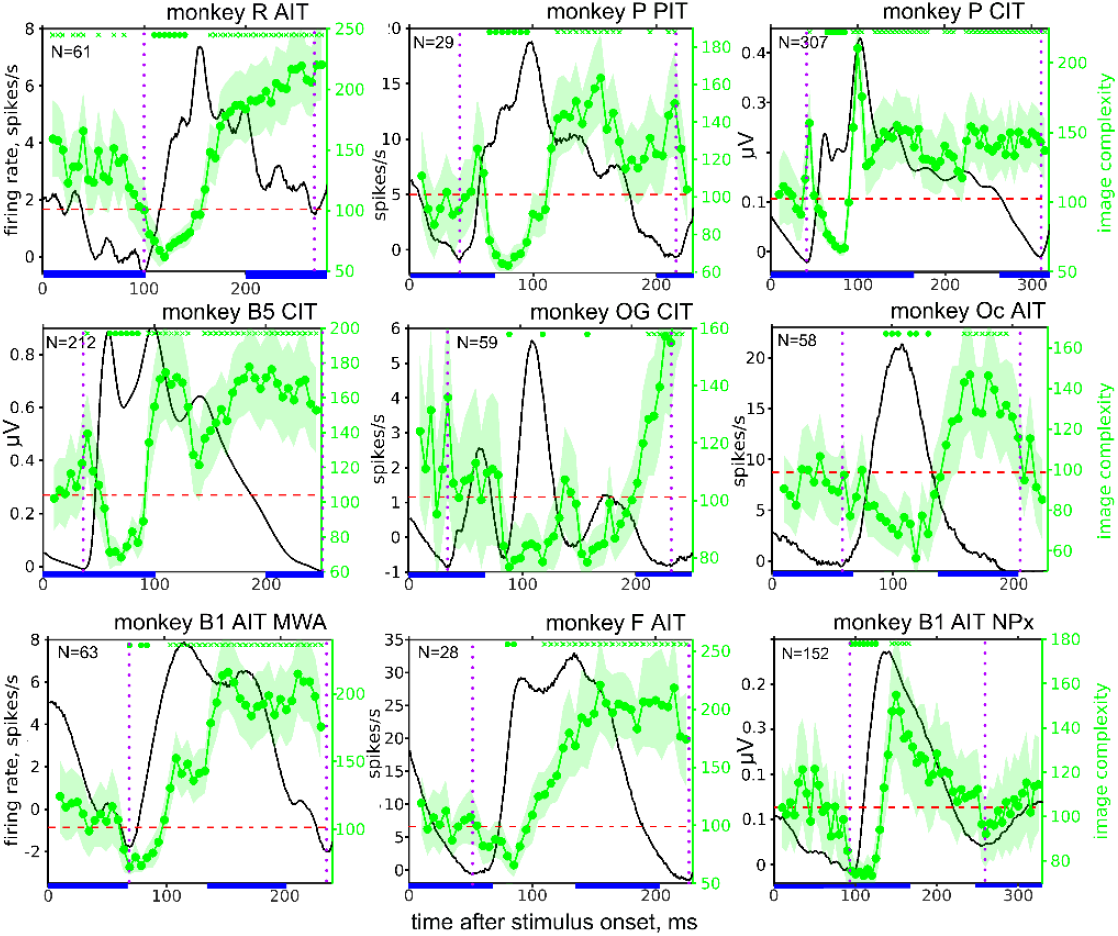
Dynamics of complexity selectivity (green traces) for 9 arrays in 7 monkeys. The average image complexity (number of parts) for the top 100 images for each 5ms time-bin, averaged over all visually responsive recording sites for each array (number of recording sites indicated at the top of each plot), are shown overlaid on the average PSTH (black traces). The average complexity of each image set is indicated by a red dashed line. The symbols at the top indicate time-bins when the preferred complexity was significantly lower (o’s) or higher (x’s) than the image set average (p < 0.05, corrected for multiple comparisons). Blue bars indicate stimulus presentation times. Dotted purple lines bracket the timepoints used in Figure 7 to interpolate the time axis.

Thus, from PIT to CIT to AIT, the earliest spikes are selective for simpler images and later spikes for more complex images, that is, images made up of more parts. This dip below followed by a rise above the image-set-average complexity was seen also in individual units (Supplementary Figure 7 top).

### Spatial frequency dynamics

Could the neuronal preference for images with increasingly more parts over time be explained by dynamics of spatial frequency tuning (Bredfeldt and Ringach 2002)? Image complexity and spatial frequency are correlated (Spearman’s ρ = 0.48 for our largest image set). We therefore analyzed the average spatial frequency of the top 100 images over the response duration. For 4 out of 9 arrays there was a dip significantly below the image-set average, and all arrays showed a gradual increase in preferred spatial frequency during the response (Supplementary Figure 8), whereas for image complexity, the dip and the gradual increase were significant in all 9 arrays (Figure 5).

To ask whether complexity or spatial frequency is the more important driver of response dynamics, we ordered the images by complexity or spatial frequency and plotted the time-course of responses to bins of images of different complexity or spatial frequency ranges. Among the top 5% of images driving the strongest response for each array, response latency increases with both complexity and spatial frequency (Supplementary Figure 9). However, if all the images are included, there is a less clear relationship between response latency and either complexity or spatial frequency. Thus, image complexity or spatial frequency alone are insufficient to explain response dynamics, and image (or feature) identity must also contribute to response dynamics.

It is generally agreed that at each stage of the early visual hierarchy, inputs encoding features from the preceding stage are combined (Hubel and Wiesel 1962; Riesenhuber and Poggio 1999). There is some evidence that the dynamics of responses in V1 and MT reiterate the hierarchical accumulation of their inputs (Ringach et al. 1997; Pack and Born 2001), in that earlier responses are less selective to orientation (V1) or to the true motion direction (MT). Thus, the earliest responses in an area seem to reflect the selectivity of antecedent areas. The fact that later parts of the response differ from earlier could be because inputs are modified by intra-areal feedback only later in the response. Early in its response a neuron may operate like an OR gate, and respond to any of its upstream inputs, but later intracortical inhibition may raise the firing threshold, making the late selectivity more like an AND operation, so the neuron responds only to conjunctions of its inputs. If this principle—the hierarchical, dynamical pooling of inputs followed by lateral feedback— continues past V1, a simple consequence could be selectivity for conjunctions of an increasing number of features in later stages and in later parts of the responses at each stage. This principle subsumes less mechanistic hypotheses that global precedes local, or category precedes individual.

### Image selectivity becomes sparser over the response duration

The canonical cortical circuit hypothesis posits pooling of feedforward excitation from antecedent areas and local feedback (Ozeki et al. 2009), which serves to normalize activity and narrow selectivity. This hypothesis predicts that early activity should be more broadly tuned as well as spread more broadly across neurons in an area, while late activity should be more narrowly tuned (in both senses) as local feedback suppresses all but the most strongly driven neurons. We therefore asked whether image selectivity becomes narrower over the duration of the response. Tuning throughout IT indeed sharpens over the duration of the response: Rank-order array-average responses (normalized per time-bin) (Figure 6) rise more steeply for late time-bins (red) than for early time-bins (blues) indicating that in later time-bins fewer images elicit the strongest responses. The same was true for individual recording sites in all the arrays; rank order responses over time for a few individual neurons from the array in monkey B1 AIT are shown in Supplementary Figure 10. Thus, tuning narrows as selectivity shifts from simple to more complex images over the duration of IT responses.

**Figure 6.**
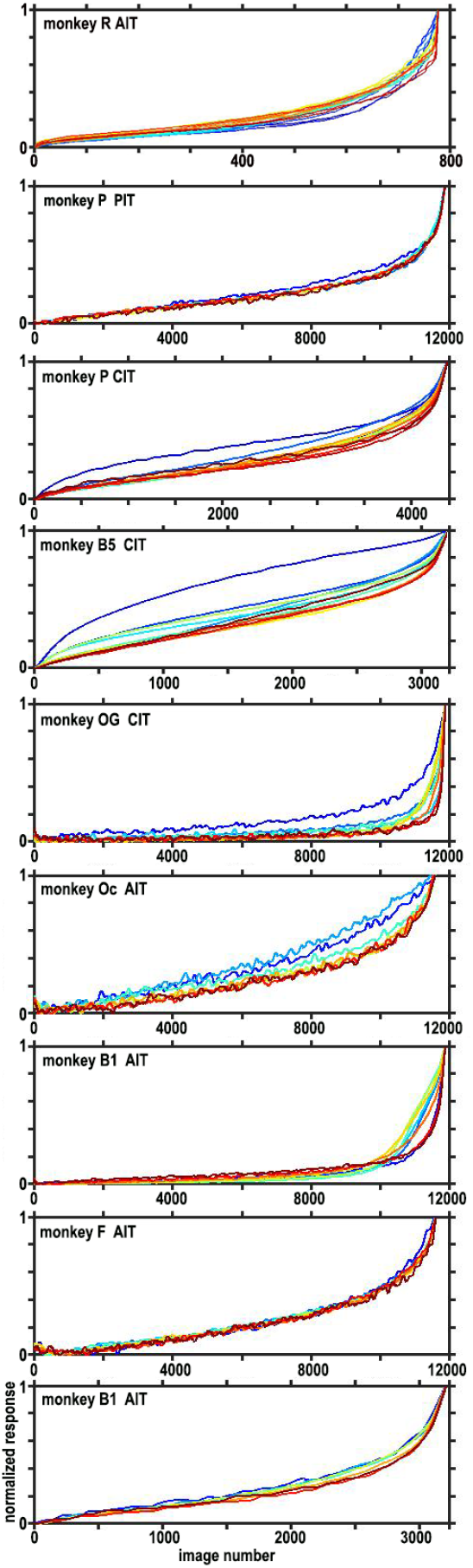
Dynamics of rank-order selectivity for all 9 focal arrays. Responses to all images in even trials were ranked by image-response order on odd trials, and vice versa, then averaged and normalized for each 20 ms time bin (color coded as in Figure 2). For all the arrays, responses became more narrowly tuned, or sparser, (responding strongly to fewer images) over the duration of the response. That is, the rank-order response curves rise more steeply in later time bins (reds) than earlier time bins (blues). Time bins color coded as in Figures 2&3.

The sparsity of image responses, also termed ‘lifetime sparseness’ (Willmore and Tolhurst 2001), has been quantified as (Treves and Rolls 1991):

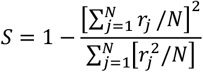

Where r_j_ is the response of a neuron to the j^th^ *image* out of a set of N images. We calculated the image sparsity for each recording site and each 10 ms time-bin starting at the beginning of the response. We selected the top 5% of the images for this quantification because the tuning sparsity (Figure 6) was most apparent among the top images, whereas the vast majority of images elicited little response yet heavily influenced the sparsity metric. The top images were determined for each site and each time-bin (cross-validated by ranking even-trial responses by odd-trial ranks, and vice versa). Then, the responses were robustly normalized among the top images before being used in the formula for sparsity. Supplementary Figure 11 shows the array-average image-sparsity for each array. For most arrays, the image-sparsity is higher in the late response compared to the early response without simply mirroring the firing rate. Similar results were obtained by quantifying image sparsity using the Gini coefficient, which directly relates to the area under the image rank response curve (Dorfman 1979).

### Increasing neuronal population sparsity over the response duration

The increasing image sparsity over the duration of the response indicates that later parts of a neuron’s response are more narrowly selective across images than earlier parts of the response. This is consistent with the idea that lateral connections eventually inhibit all but the strongest responses. If this response narrowing in IT neurons is due to intracortical suppression, as has been proposed for V1 (Ringach et al. 2002; Ringach et al. 2003), we would also expect the population of responding neurons to narrow over time. That is, we expect that initially many neurons respond to the same stimulus, then these simultaneously activated neurons inhibit each other, thus reducing the number of strongly responsive neurons. To ask whether the number of neurons responding to each image also decreases over the response, we measured an analogous population sparsity (Willmore and Tolhurst 2001), per time bin, averaged across images: 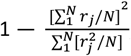, where r is now the response to each image of the jth *neuron* out of N visually responsive neurons in each array. The neuronal population sparsity is calculated per image and then averaged over responses to the top 5% of images. We again considered only the top images because only images producing strong responses are expected to recruit intracortical inhibition. Neuronal population sparsity for most of the arrays also peaks late in the response (Supplementary Figure 12). Thus, the number of neurons responding strongly to each image decreases over the duration of the response.

A general trend across IT Cn be revealed by interpolating all these metrics―PSTHs, complexity selectivity, spatial frequency selectivity, image sparsity, and neuronal population sparsity—onto a common, normalized time axis across arrays (Figure 7). On average, across all our IT data sets, complexity first decreases below the image-set average, then rises over the duration of the response. Spatial frequency also gradually rises over the response duration. Image sparsity and neuronal population sparsity both rise late in the response.

**Figure 7.**
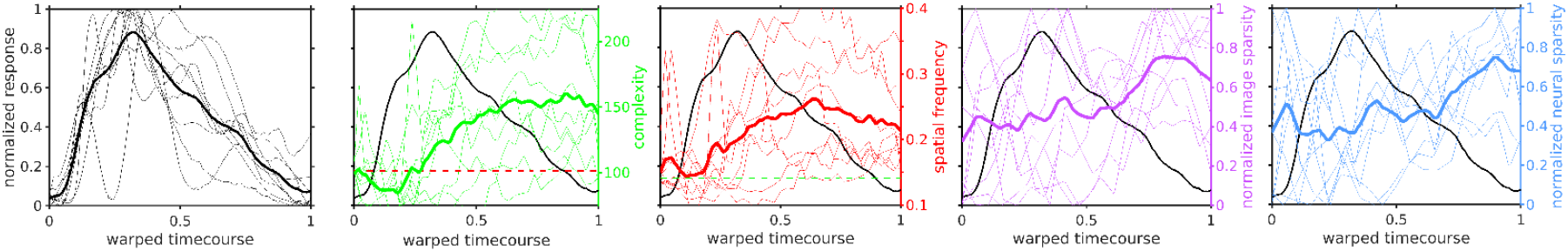
Metrics of dynamics in all 9 arrays, interpolated to a common, normalized time axis (bracketed by dotted purple lines in Figure 5) and then averaged. Thin lines indicate individual arrays; thick lines show the average over all 9 arrays. On average across the 9 focal IT arrays image selectivity shifts from simple to more complex (green), and becomes sparser (purple) over the duration of the response. The neuronal population response (blue) also becomes sparser over time.

### Dynamics of representational similarity with DNN layers throughout the visual hierarchy

Our results recording from focal IT arrays show response dynamics in which tuning shifts to increasing image complexity and spatial frequency. We propose that these dynamics in IT recapitulate antecedent processing stages along the visual hierarchy. Therefore, we asked whether selectivity would be similar between AIT and antecedent areas, especially early in the response period, and whether selectivity similarity to AIT would occur early in the response for early visual areas. We implanted, chronically, four 45mm long Neuropixels arrays that spanned from V1, through V2, V3, V4, to anterior IT (Figure 8 left, shows these arrays visualized in CT scans, with the probe trajectory aligned to the brain anatomy). Recording simultaneously across the ventral stream from V1 to AIT, we calculated representational similarity between AIT and two different sections of the probe representing earlier and later visual areas (Fig. S13). There was significant representation similarity between areas. Two data sets showed earlier correlations between AIT and V2/V3 compared to AIT and CIT (monkey B1) and between AIT and V1/V2 compared to AIT and PIT (monkey V). However there was no clear timing difference between the correlations for AIT and PIT compared to AIT and V1 in monkey P, and for monkey T, the correlations between AIT and CIT were earlier than the correlations between AIT and V2/V3. The inconclusive timing analysis may be because the recorded sites across areas did not necessarily have overlapping receptive fields or belong to the same selectivity network {Bao, 2020 #6380}.

**Figure 8.**
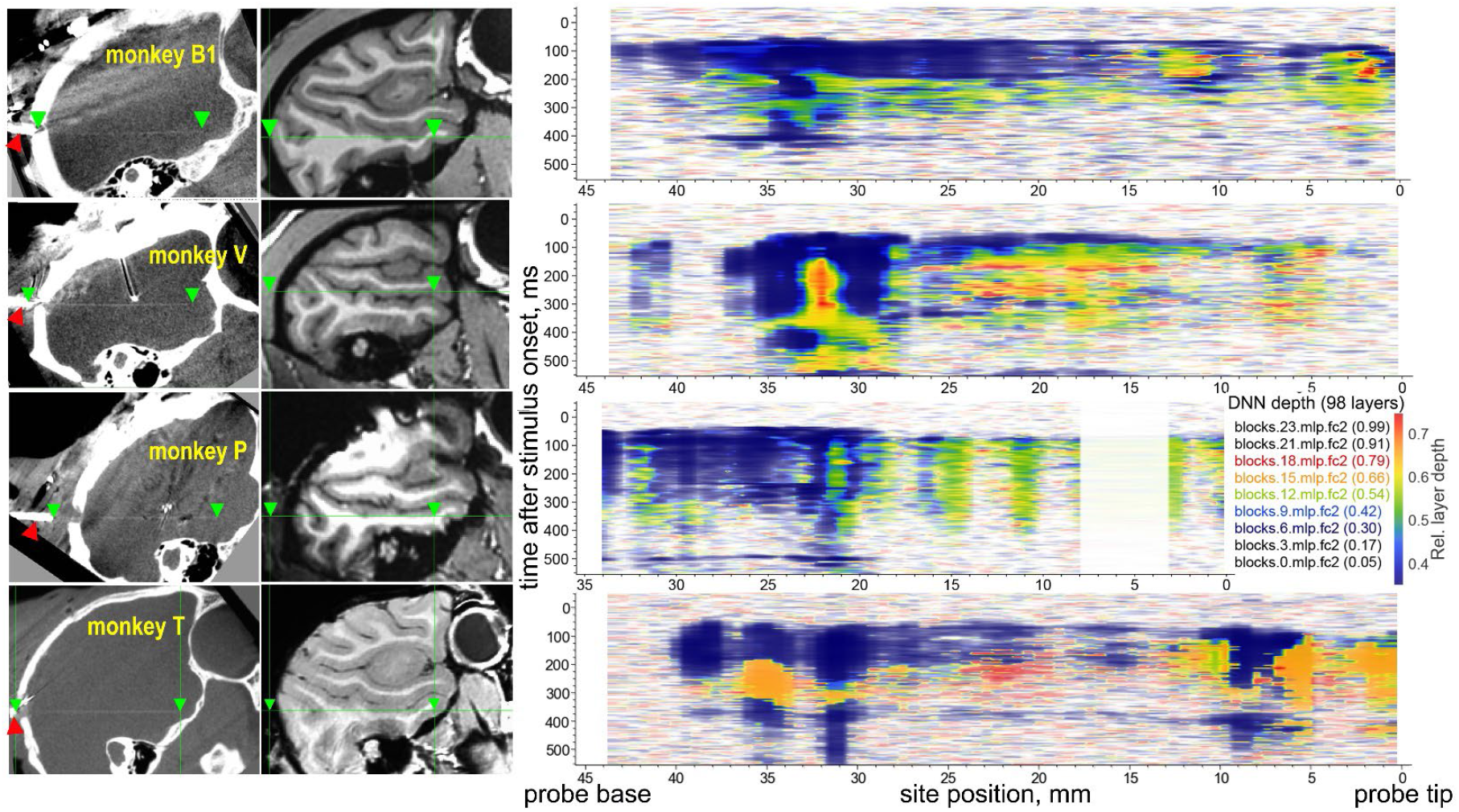
Selectivity dynamics along the visual hierarchy in four monkeys with chronic Neuropixels probes spanning from V1 to AIT. (left) The four probes visualized by CT scans. Each probe is visible as a faint horizontal line in sagittal CT sections. The green arrowheads indicate the ends of each probe (45mm apart), and the red arrowhead points to the base electronics of the probe. The linear density in monkey V’s CT above the Neuropixels array is a microwire array. The white dot densities above monkey P’s Neuropixels array correspond to an FMA (monkey P PIT array). The second column shows an anatomical MRI aligned to the CT, with the Neuropixels probe span indicated by green arrowheads. On the right, the heatmaps indicate for each location on the probe and each 5ms time-bin, the depth of the most similar layer in ViT by representation similarity analysis. The color saturation indicates the magnitude of representation similarity normalized per heatmap. The gap at 3 - 8 mm in the heatmap for monkey P indicates electrodes excluded from the recording because they were not visually responsive in a probe-wide survey.

Therefore, instead of directly comparing different areas, we analyzed representation similarity to DNNs as a proxy. IT selectivity dynamics are reflected by a shift over time in DNN representational similarity from shallow to deep layers (Figure 4 bottom row). We asked whether processing stages along the visual hierarchy show representational similarity to progressively deeper DNN layers, both over the response duration and over the processing hierarchy. The depth of the DNN layer in ViT with the highest representation similarity increases (blue → red) both with time during the response (y axis) and with position along the visual hierarchy (x-axis, V1 to AIT from left to right). Thus, the most representationally similar DNN layers are shallower for early visual areas and deeper for later visual areas, and shallower for early epochs of the response and deeper for later epochs.

## Discussion

We recorded from 13 chronically implanted multi-site arrays in IT of 9 monkeys. The chronic arrays allowed concatenation of data over days to weeks, thus allowing us to analyze response dynamics to very large image sets. By inspection, the earliest selectivity in face-selective regions in IT was for simple images—large, round, often warm-colored objects, including large clear faces, but the selectivity shifted over the duration of the response to more complex images, including faces and partly occluded faces, and finally, towards the end of the response, to whole individuals. A shift from simple images (few parts/low spatial frequency) to more complex images (more parts/high spatial frequency) was seen in both face-selective and non-face-selective arrays and in individual sites in each array. We suggest that these selectivity dynamics reflect sequential processing stages upstream to the cells recorded: the earliest responses in each area resemble the inputs before lateral suppression modifies the responses, so traces of the response properties of earlier input areas appear transiently in higher areas. Thus, we propose, the response dynamics in higher areas recapitulate antecedents in the visual hierarchy.

Preferred image complexity and preferred image spatial frequency both increased during the response. We could not rule out either one as the dominant parameter underlying the selectivity dynamics, although it seems less likely that the visual system is a Fourier analyzer of images than that the visual system encodes images by sequential conjunction of features. Several lines of evidence support the hypothesis that the visual hierarchy combines image features, rather than analyzing spatial frequency: 1) contours are elemental image features encoded in V1 (Hubel and Wiesel 1962); 2) corner and curvature selectivity in V2 (Hubel and Wiesel 1968; Hubel and Livingstone 1987) can be constructed by combining two or more contours; 3) the number of contours encoded in PIT increases over the response duration (Brincat and Connor 2006); and 4) some face-selective neurons in PIT encode specifically the image feature of the contralateral eye (Issa and DiCarlo 2012). Since images with more parts tend to have higher spatial frequency, selectivity for image parts should be correlated, perhaps with varying consistency, with spatial frequency.

Previous studies on the dynamics of macaque IT selectivity have used smaller image sets designed to test specific hypotheses. Sugase et al (1999) recorded from macaque IT using 38 images of faces and objects, and reported that information about image category (face vs object) emerged early and transiently, whereas information about face identity and expression emerged later and was more sustained; they interpreted this result as indicating that global information is used as a ‘header’ to prepare destination areas for receiving more detailed information. In a previous study on face cells (Tsao et al. 2006) we used 96 images of human faces, bodies, fruits, gadgets, hands and scrambled images (16 each) and reported that information about whether an image is a face or not emerges earlier than information about face identity. We interpreted this as indicating that face categorization is distinct from face analysis, with the initial categorization step involving holistic non-linear calculations, and subsequent individuation or identification computations being linear (Tsao and Livingstone 2008). Koyano et al. (2021) found in macaque IT face patches a delayed suppression to average faces leading to a delayed preferred response to more distinct faces. These prior studies posit task-, or area-specific computations. The first two hypotheses above, 1) that some kind of ‘header’ information prepares destination areas for more detailed information or 2) that face detection is qualitatively different from face analysis, both require special circuitry with task-specific functionalities that change over the duration of the response.

Our results are consistent with these prior studies, but they support a more universal hypothesis: the computation of pooling followed by intracortical inhibition at each stage of the visual hierarchy (Riesenhuber and Poggio 2000). This principle is consistent with the proposal of a canonical cortical circuit performing similar computations on whatever inputs it gets (Douglas et al. 1989; Riesenhuber and Poggio 1999). Our results provide further evidence to support Tamura and Tanaka (2001), who found that IT responses become more selective, responding to fewer images, over the duration of the response; they interpreted this as indicating that among neurons initially activated, weakly responding neurons are suppressed by mutual lateral inhibition, so only strong responses persist. Consistent with this idea of intracortical inhibition narrowing IT selectivity, local application of the GABA antagonist bicuculline increases neuronal responsiveness in IT to previously poorly effective images (Wang et al. 2000).

Prior studies showing categorization preceding individuation, average preceding distinct, or global information preceding local, can be interpreted as manifestations of the principle of a canonical cortical circuit that combines increasingly more features at each stage: Individuation could require conjunction of more features than categorization; distinct face processing could require more features than identifying an average face; local information comprises more features than global; and, to circle back to the observation that motivated this study (Figure 1), whole individuals have more parts than faces do. That we observe similar dynamics for both face-selective arrays and non-face-selective arrays is consistent with similar computations across cortex, not as qualitatively distinct task-specific computations.

Riesenhuber & Poggio (2000) modeled the visual hierarchy as a series of pooling operations, each followed by a maximum operation, where mutual inhibition leads to a winner-take-all output. They envisioned pooling and maximum operations as occurring simultaneously, such that the output at each stage is a discrete function of both operations, and there would be no reiteration of earlier selectivities at each stage. But in posterior IT, responses to individual contours (parts) are faster and more transient than responses to multiple-contour stimuli (Brincat and Connor 2006), suggesting that outputs of earlier visual areas, like selectivity for individual contours in V1, bleed into the early responses of subsequent areas, like PIT. Furthermore, the dynamic behavior observed in many studies (the transiency of responses, shifts from local to global or categorical to individual, shifts from single to multi-part tuning, and our results here) suggests a delay at some stage(s) between pooling operations and intracortical suppression, such that the earliest spikes in a given area are less refined by intracortical suppression than later spikes. This would make sense, because even if synaptic delays are shorter than the time-scales considered here, intracortical suppression at each stage could plausibly take some time to build up, particularly from remote or less activated neighbors. Indeed, if there is such a delay at multiple stages we might expect a cascade of response properties reflecting first pooling at each stage, then only later the effects of intracortical interactions at each stage, consistent with the dynamic changes in selectivity observed here. We were unable to show directly that correlations between areas show graded dynamics, but we found that correlations between DNN layers and IT did show graded dynamics. To extrapolate further, we suggest that, at each stage, a neuron’s baseline firing threshold is relatively low, such that early in its response the neuron operates like an OR gate, reflecting its upstream inputs, but later intracortical inhibition raises the effective firing threshold, making the late selectivity more like an AND operation, thereby creating new, more complex, selectivities.

The results of all these prior studies are consistent with our results, but our results further show that the early broad selectivity is to images of low complexity, and over the duration of the response, IT neurons became more selective for more complex images, defined simply as images with more parts. We envision the pooling at each stage to encode combinations of features from the preceding stage of the hierarchy, resulting in each stage representing objects as conjunctions of increasing number of parts. Concurrently the image selectivity narrowed, so many images elicited strong responses at short latencies, but fewer images elicited strong responses at longer latencies. The similarity of the selectivity dynamics between highly face-selective arrays and non-face-selective arrays support the idea of a canonical circuit, performing similar computations, rather than task-specific circuitry, even for face-selective domains.

How can we explain the exquisite selectivity of entire domains specifically responsive to, for example, faces, by a generic architecture? Clusters of highly face-selective neurons in humans and monkeys have been interpreted as evidence for the evolution of circuitry innately specialized for detection and analysis of faces in particular (Kanwisher 2000). We have argued (Arcaro and Livingstone 2024) instead that self-organizing Hebbian mechanisms and extensive experience seeing faces is sufficient to account for face-selective domains, particularly since seeing faces early in life is essential for the development of face domains (Arcaro et al. 2017). Exquisitely specific biological functionalities, like eyes, were once considered evidence against evolution (Darwin 1859). The plausibility of highly selective domains emerging in a domain-general architecture is supported by studies on neural networks: category-specific nodes emerge in deep neural networks (with a generic, modular, architecture) trained only on individuation, not categorization (Dobs et al. 2022; Prince et al. 2024), and clusters of category-specific nodes, or domains, emerge in generic-architecture networks that incorporate topological constraints (Blauch et al. 2022; Margalit et al. 2024).

Lastly, our results suggest that response dynamics reflect a cascade of pooling followed by normalization operations occurring at multiple hierarchical stages, rather than the more teleological idea of a progression towards the optimal stimulus for a given neuron. That is, we do not think these dynamics represent some kind of temporal code carrying changing information over the response duration, but rather a reflection of the hierarchical circuitry of the visual pathway.

## Acknowledgments

This work was supported by NIH grants P30 EY012196, R01 EY025670, and R01 NS123778 to MSL.

## Notes

### Competing Interest Statement

The authors have declared no competing interest.

