## Supplementary Figures for "Response dynamics in macaque ventral stream recapitulate the visual hierarchy"

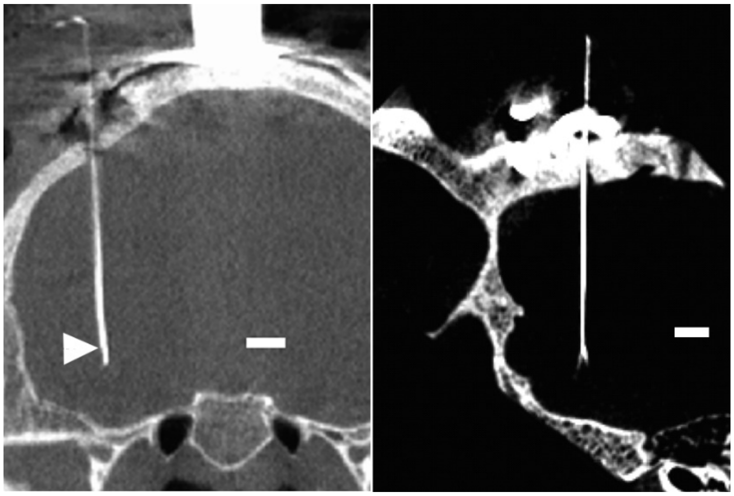

**Supplementary Figure 1.** High resolution CT scan of the smallest (left) and largest (right) splay of the Microwire Arrays used here. The bare wires are pushed out of the guide tube 3-5mm. The junction between the guide tube and the bare wires is visible as a slight offset at the arrowhead on the left and a splaying out on the right; the wires splay out less than 1 mm in the monkey on the left (monkey R AIT), and less than 2mm in the monkey on the right (monkey B1 AIT). Scale bars indicate 5mm.

**Supplementary Table 1.** Array type, location, number of images, and mean repetitions per image per set of chronic recording sessions.

| Monkey | Location | Array type | # unique images | Mean # images/ category | Mean repetitions per image | Stimulus duration (ms) | Inter-stimulus interval (ms) |
|---|---|---|---|---|---|---|---|
| monkey R | AIT | Microwire array | 776 | 4 | 33.6 | 200 | 200 |
| monkey R | AIT | Microwire array | 8875 | 15.6 | 37.7 | 100 | 100 |
| monkey P | PIT | Floating Microelectrode Array | 8865 | 15.6 | 40.0 | 67 | 134 |
| monkey P | CIT | 10mm Neuropixels | 4216 | 7.4 | 36.2 | 167 | 100 |
| monkey B5 | CIT | 10mm Neuropixels | 3178 | 7.4 | 58.1 | 167 | 100 |
| monkey OG | CIT | Microwire array | 8853 | 15.6 | 59.4 | 67 | 134 |
| Monkey Oc | AIT | Microwire array | 8871 | 15.6 | 42.3 | 67 | 67 |
| Monkey B1 | AIT | Microwire array | 8877 | 15.6 | 58.8 | 67 | 67 |
| Monkey F | AIT | Microwire array | 8875 | 15.6 | 42.6 | 67 | 67 |
| Monkey B1 | AIT | 45mm Neuropixels | 3178 | 7.4 | 19.7 | 200 | 100 |
| Monkey B1 | V1 to AIT | 45mm Neuropixels | 3178 | 7.4 | 9.1 | 100 | 200 |
| Monkey V | V1 to AIT | 45mm Neuropixels | 3178 | 7.4 | 14.2 | 200 | 267 |
| Monkey P | V1 to AIT | 45mm Neuropixels | 3178 | 7.4 | 15.8 | 200 | 234 |
| Monkey T | V1 to AIT | 45mm Neuropixels | 776 | 4 | 32.4 | 100 | 167 |

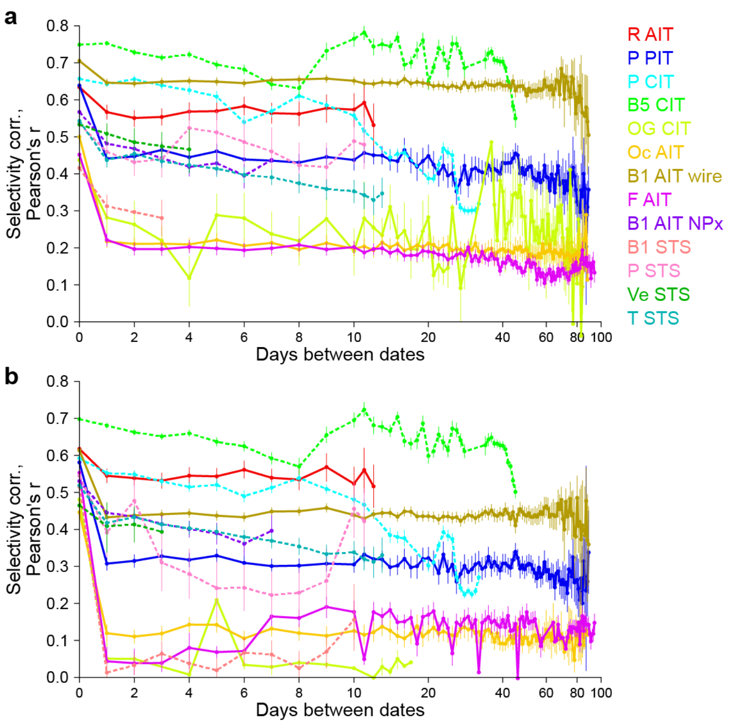


**Supplementary Figure 2.** Stability of response selectivity over chronic recording sessions. (a) Image-category level selectivity stability over recording sessions for the data used in this study. Each day's pattern of category selectivity was correlated with that in every other day of recording. Within-day correlation (0 days between dates) was quantified as trial-split-half self-consistency. (b) Individual-image level selectivity stability over sessions. Points and error bars indicate mean and bootstrap 95% confidence interval across all neurons and session pairs with the same days between dates. Dashed lines correspond to Neuropixels recordings; solid lines correspond to other recordings (MWAs and one FMA). The two subplots show different date ranges for datasets B1 ST S and OG CIT because different sets of neurons reached the threshold for being considered visually selective, which was defined by selectivity among either image categories (a) or individual images (b).

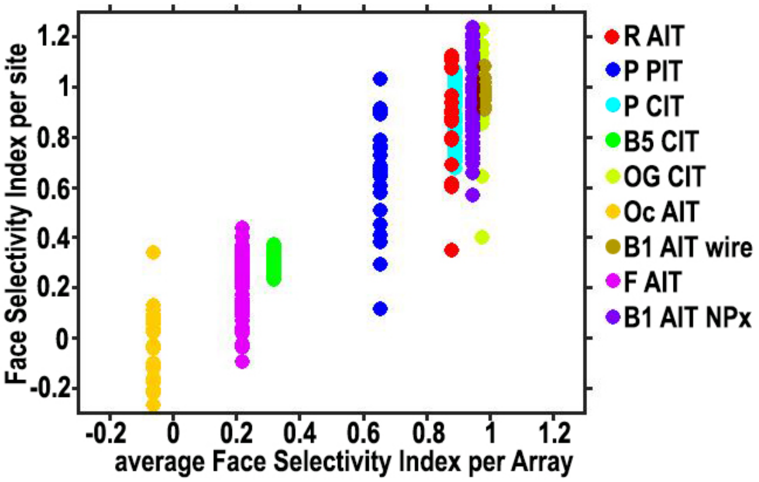


**Supplementary Figure 3.** Face Selectivity Index for all the focal arrays in this study (i.e. excluding the NPx arrays that spanned visual areas in Figure 7). X-axis is the average Face Selectivity Index for each array; Y-axis is the Face Selectivity Index for each recording site in each array. Each color represents a different array, as indicated. Face Selectivity Index= (response-to-faces minus response-to-objects) / (response-to-faces + response-to-objects).

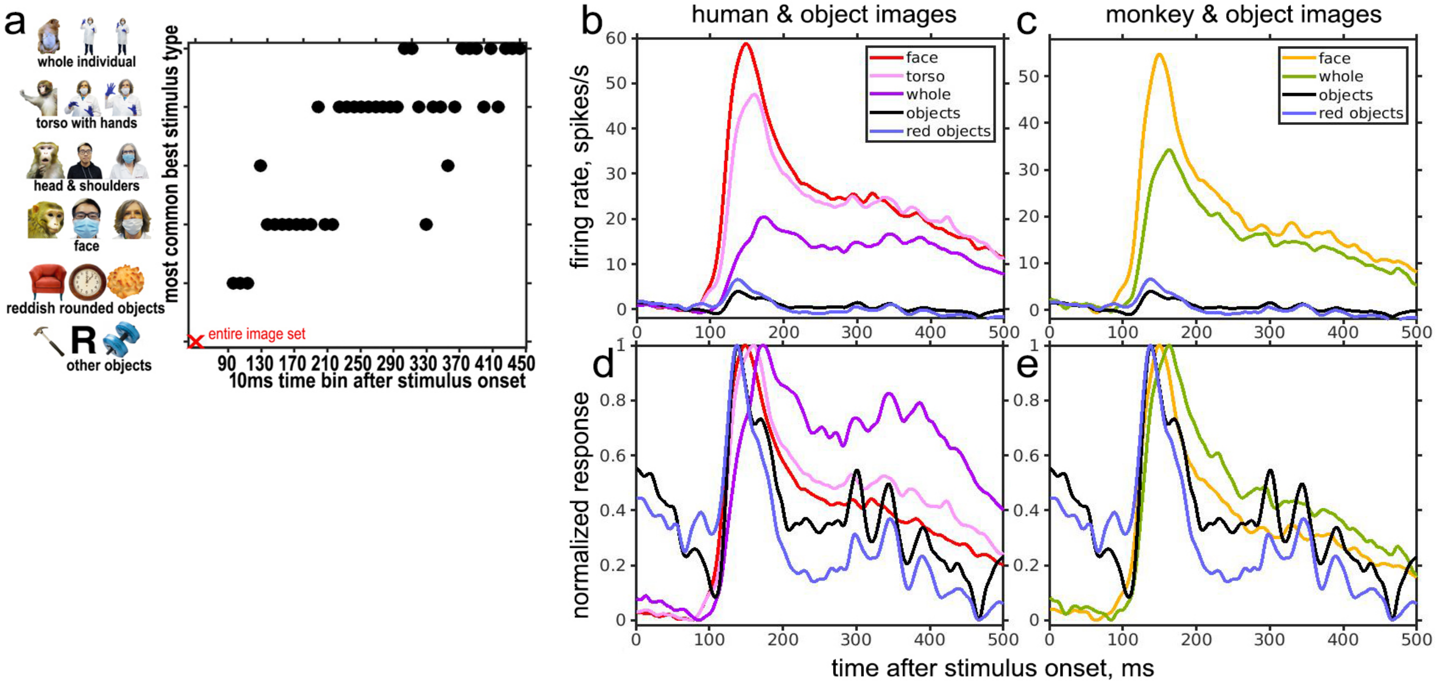


**Supplementary Figure 4.** Response dynamics for different image categories. (a) The most common top-image category for each of the 31 recording sites in the microwire array in monkey R AIT in each 10ms time-bin after stimulus onset. The red 'x' indicates that the most common category in the full image set was inanimate objects that were not reddish and rounded. Examples of each category are shown to the left. (b) Array-average PSTHs in response to human faces, human torsos, whole humans, reddish round objects, and other objects. (c) Array-average PSTHs in response to monkey faces, whole monkeys, reddish rounded objects, and other objects. (d & e) The array-average PSTHs normalized per category, highlighting differences in the time-courses. All humans pictured are authors of this study.

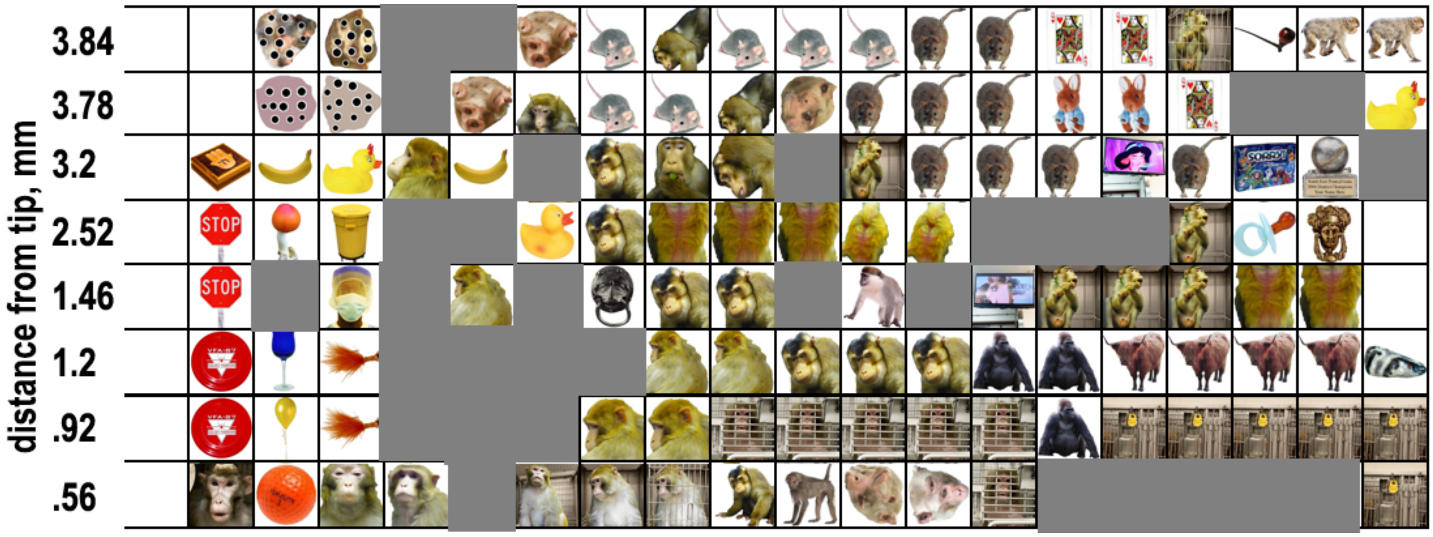


**Supplementary Figure 5.** Magnified images for 8 sites from Figure 2, at the indicated distances from the probe tip. Images of humans have been grayed out.

Identity of the top images for each site, L to R:

3.84: scrambled monkey face with black dots, same, human face, same, scrambled monkey face, mouse, monkey facing downwards, mouse, same, same, backside of a rat, same, queen of hearts, same, monkey in a cage, pipe, monkey, same.

3.78: dots on a pink background, dots on a tan background, human face, scrambled monkey face, monkey torso, mouse, mouse, monkey facing downwards, scrambled monkey face, rat backside, same, same, toy rabbit, same, queen of hearts, human torso, same, rubber duck.

3.2: box, banana, rubber duck, monkey profile, banana, baseball player, monkey face, same, same, baseball player, monkey in cage, rat backside, same, same, female cartoon face on monitor, rat backside, game box, trophy, human face in helmet.

2.52: stop sign, mushroom, yellow trash can, human torso, human face, rubber duck, monkey face, monkey butt, same, same, another monkey butt, same, baseball player, same, same, monkey in cage, pacifier, door knocker.
1.46: stop sign, human face, human face in PPE, human face, monkey torso, human torso with PPE, door knocker, monkey face, same, baseball player, monkey, human face, cartoon animal face on monitor, monkey in cage, same, same, monkey butt, same.
1.2: red frisbee, wine glass, fishing lure, human face, human face with PPE, human torso with PPE, same, monkey torso, same, monkey face, same, same, gorilla, same, water buffalo, same, same, same, same, rock.
0.92: red frisbee, balloon, fishing lure, human face, human face, human torso with PPE, monkey face, same, monkey face inside cage, same, same, same, same, gorilla, water bottle and cage tag on cage, same, same, same, same.
0.56: monkey face, orange golf ball, monkey face, monkey face, human torso with PPE, monkey in cage, monkey in cage, same, monkey, monkey, scrambled monkey face, scrambled monkey face, monkey face in cage, baseball card, same, same, same, same, water bottle and cage tag on cage.
.

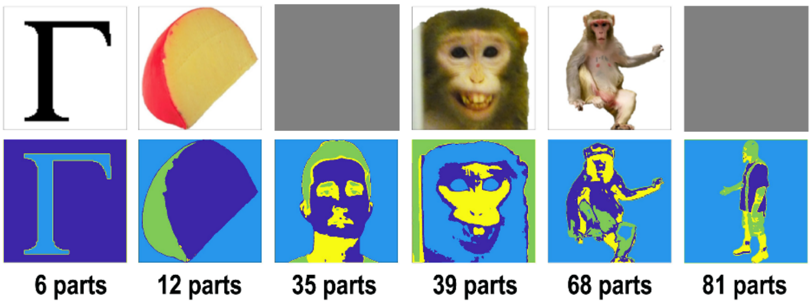


**Supplementary Figure 6.** Segmentation of images into parts. Images are first segmented into four color clusters found by K-means clustering. The color-labeled partitions are then separated into non-contiguous parts by connected-components labeling. Human images have been grayed out.

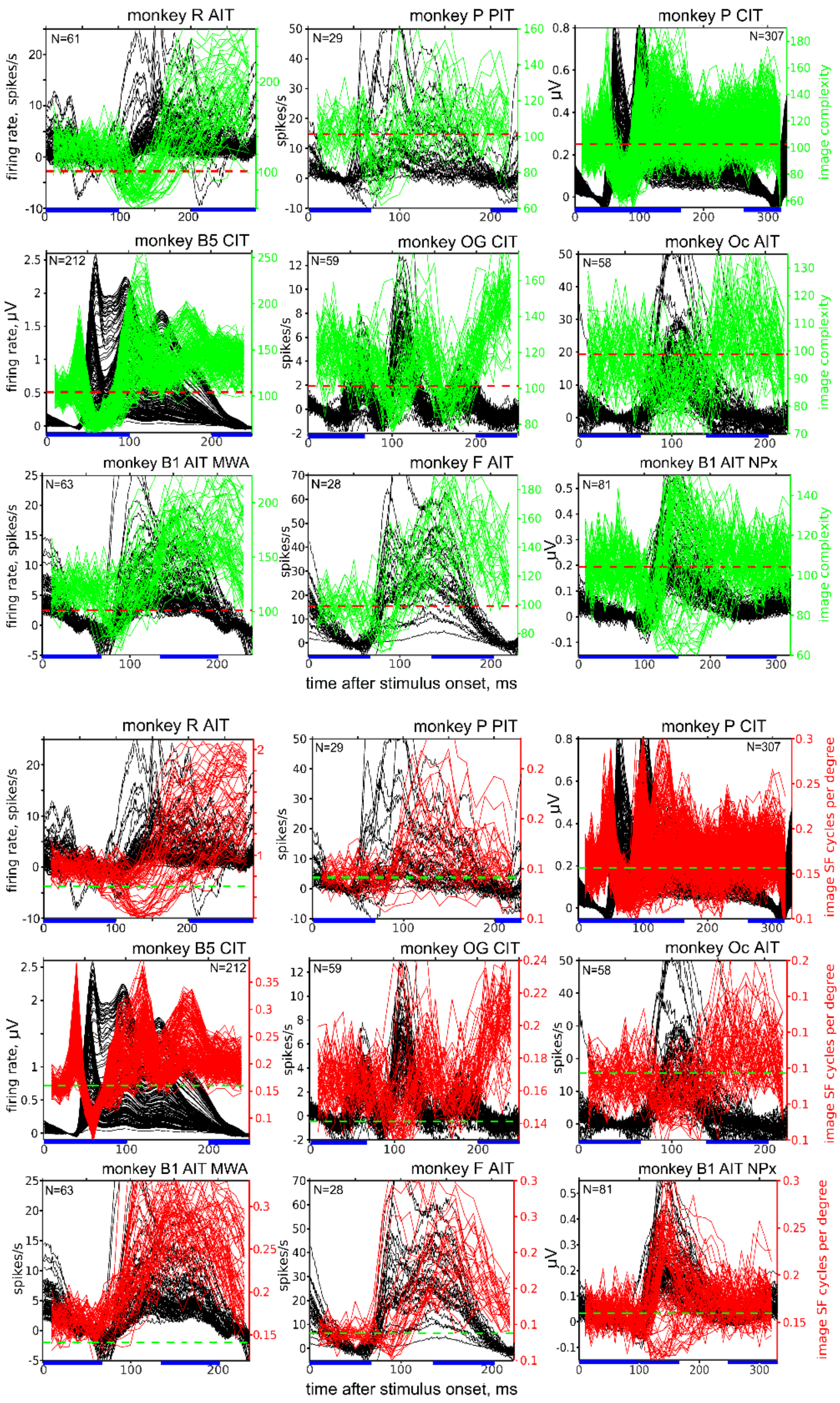


**Supplementary Figure 7.** Image complexity (green traces, top three rows) and image spatial frequency (red traces, bottom three rows) averaged over the top 100 images for each 10ms time-bin for each visually responsive recording site in each array. Black traces show the PSTH for each site. The dashed lines indicate the average image complexity (dashed red lines) or the average image spatial frequency (dashed green lines) of each image set.

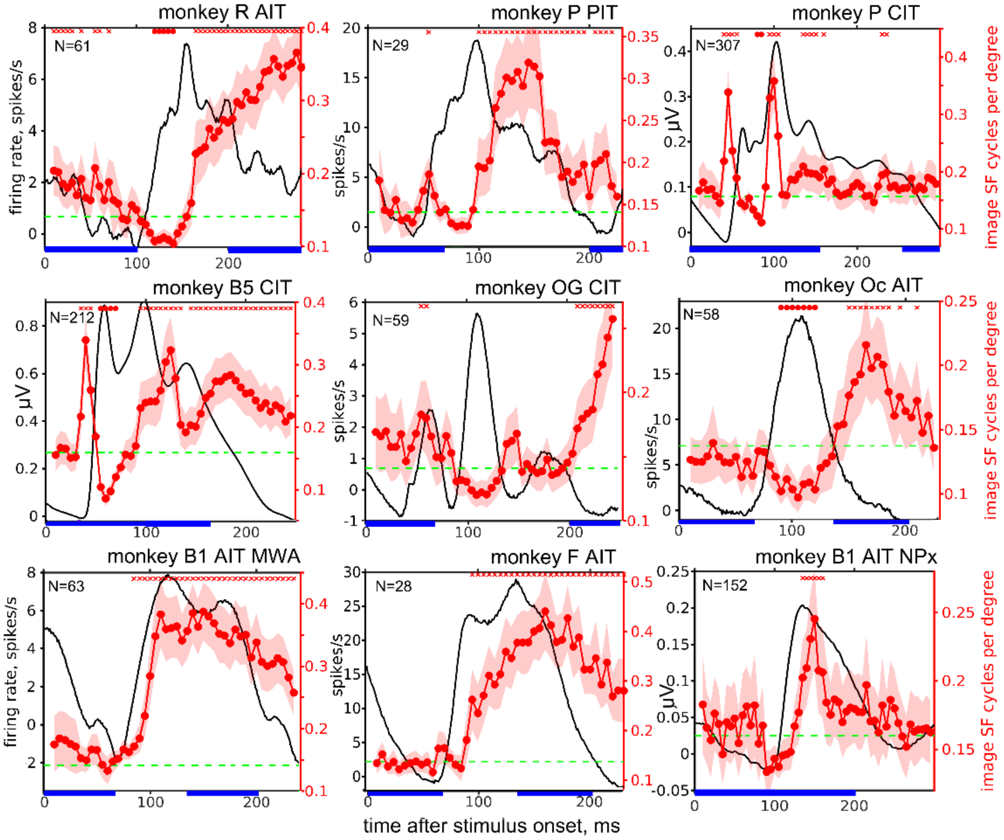


**Supplementary Figure 8.** Dynamics of spatial frequency selectivity for 9 arrays in 7 monkeys. The average image spatial frequency for the top 100 images for each 5ms time-bin are plotted in red, and the average PSTH is shown in black. The average spatial frequency of each image set is indicated by a dashed green line. The red symbols at the top of each plot indicate time-bins when the spatial frequency of the top 100 images was significantly smaller (o's) or larger (x's) than the average spatial frequency of the image set ( p<0.05, corrected for multiple comparisons).

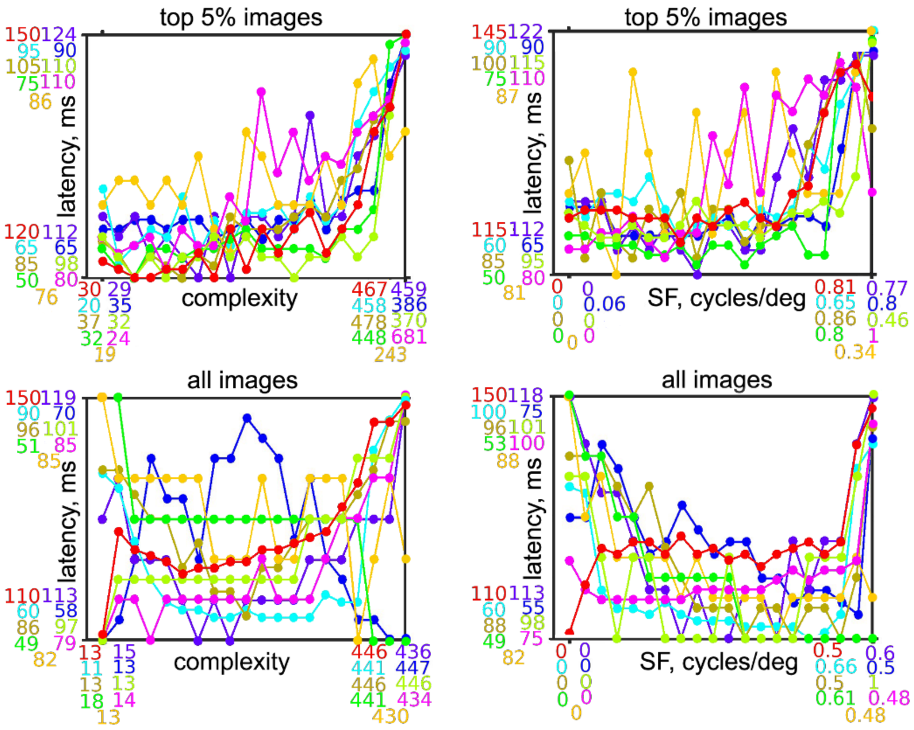


**Supplementary Figure 9.** Latency of responses to the top 5% of images (top) or all the images (bottom) as a function of complexity (left) or spatial frequency (right). The top images or the entire image set were ordered and binned into 20 bins, according to image complexity or spatial frequency, and the array average response was calculated. Then the time to half peak response was found. Each array is indicated by color, as in Figure 4 bottom. The ranges for each array for both axes are indicated by the same colors.

**Supplementary Figure 10.** Rank-order dynamics for 4 individual recording sites from the B1 AIT wire array. Responses to all images in even trials were ranked by image order on odd trials, and vice versa, then averaged and normalized per 20 ms time bin. Time bins color coded as in Figures 2&3.

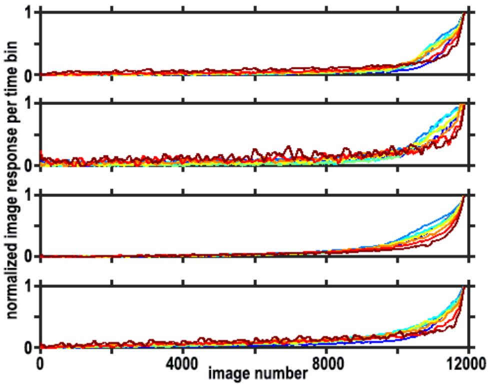


**Supplementary Figure 11.** Image sparsity averaged over all responsive sites in each focal array. The black trace (left y-axis) is the average PSTH, and the pink trace (right y-axis) is the image-sparsity for each 10ms time-bin. Shading indicates 95% confidence intervals.

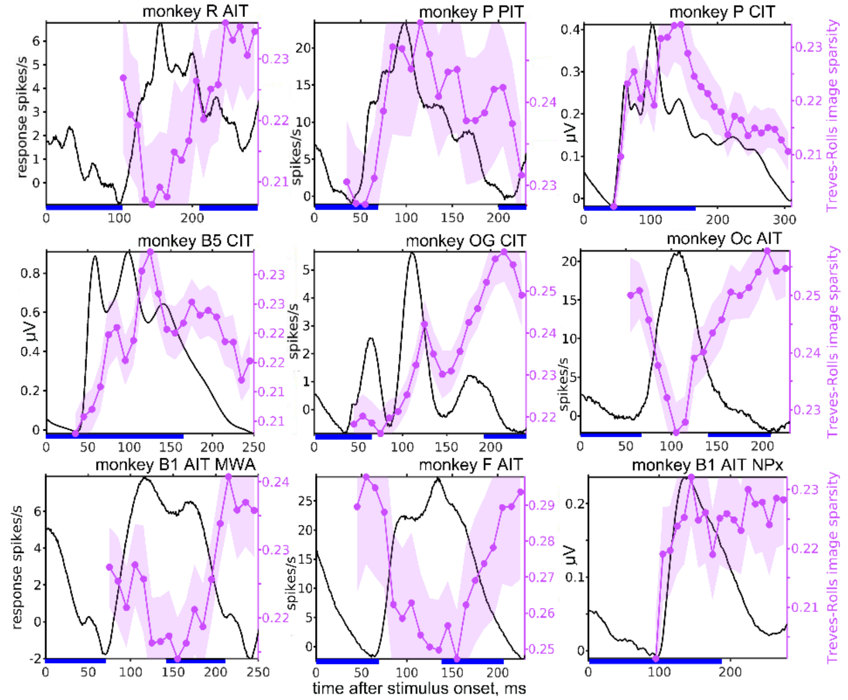


---

**Supplementary Figure 12.** Neuronal population-sparsity averaged over the sparsity to the top 5% of the images. The black trace (left y-axis) is the average PSTH, and the blue trace (right y-axis) is the neural-sparsity for each 10ms time-bin. Shading indicates 95% confidence intervals.

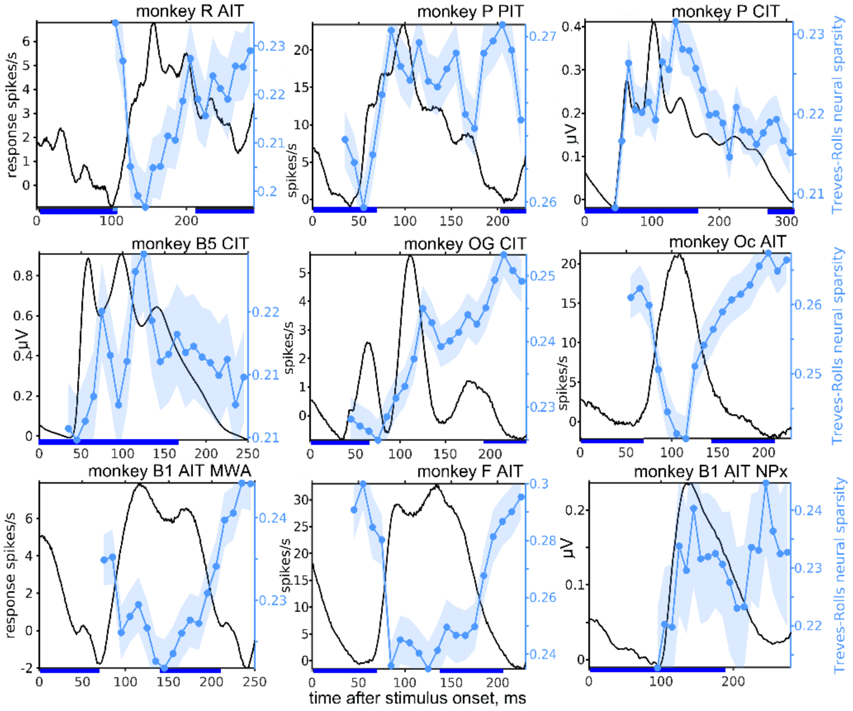


.

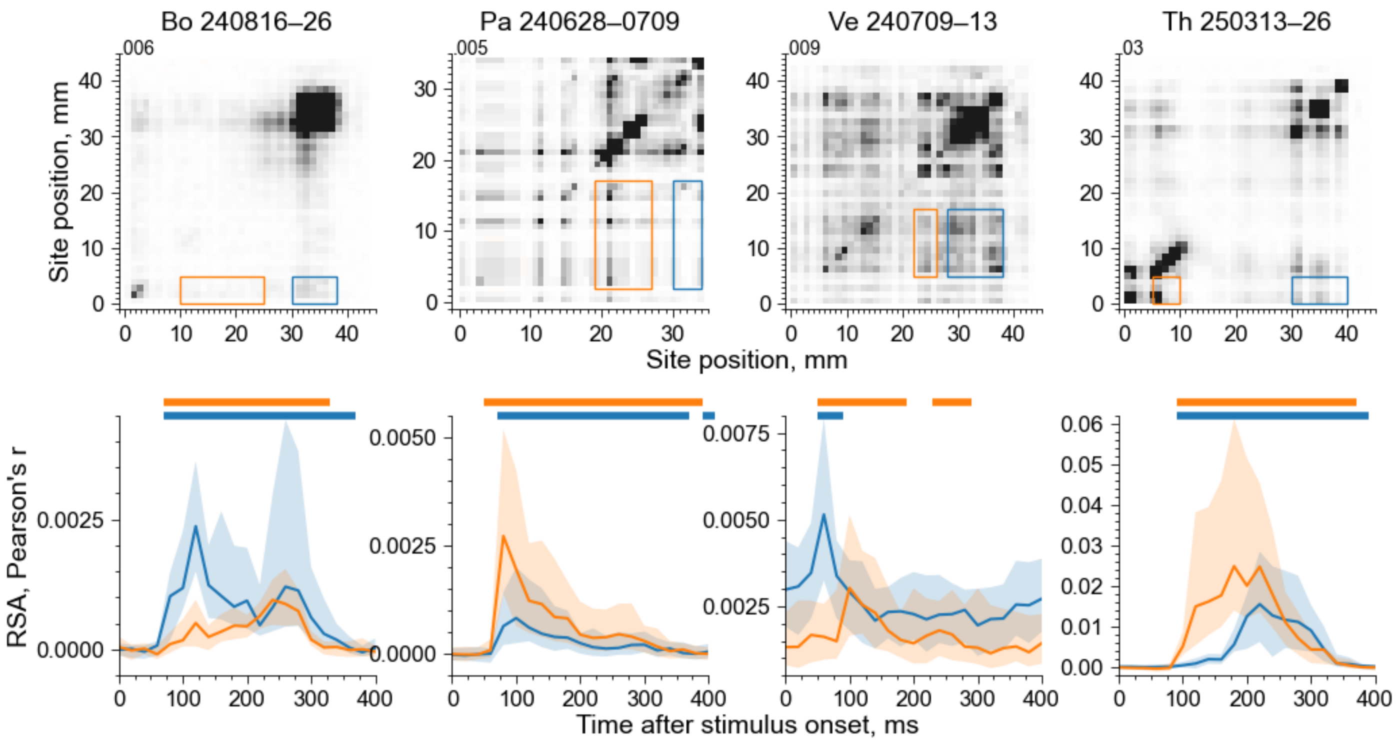


**Supplementary Figure 13.** Representational similarity between AIT responses (indicated by the vertical extent of the blue and orange boxes) and earlier visual areas (indicated by the horizontal extent of the blue and orange boxes). (top) RSA correlations between pairs of groups of ten adjacent sites. Darker = stronger correlations; black color corresponds to the 97.5th-percentile correlation, indicated by the number at the top of the y axis in each plot. Site positions in mm are relative to the probe tip (0 mm). (bottom) Time-course of inter-areal correlations between activity near the probe tip, AIT, and earlier visual areas as indicated in the top row. Each line and shading indicate the median and inter-quartile ranges across pairs of site groups. Horizontal bars above indicate time bins where the population RSA is statistically different from baseline values (mean between 0–50 ms) ($p < 0.01$, one-sample, one-tailed Wilcoxon signed-rank test, FDR-corrected).
